# A population of bang-bang switches of defective interfering particles makes within-host dynamics of dengue virus controllable

**DOI:** 10.1101/479527

**Authors:** Tarunendu Mapder, Sam Clifford, John Aaskov, Kevin Burrage

## Abstract

The titre of virus in a dengue patient and the duration of this viraemia has a profound effect on whether or not a mosquito will become infected when it feeds on the patient and this, in turn is a key driver of the magnitude of a dengue outbreak. As mosquitoes require 100-1000 times more virus to become infected than a patient, the transmission of dengue virus from a patient to a mosquito is a vulnerability that may be able to be targeted to improve disease control. The intrinsic variability in the within-host dynamics of viraemias is explored for a population of patients using the method of population of models (POMs). A dataset from 207 patients is used to calibrate 20,000 models for the infection kinetics for each of the four dengue virus serotypes. The effect of adding defective dengue virus interfering particles to patients as a therapeutic is evaluated using the calibrated POMs in a bang-bang optimal control setting.

**Author summary:** Dengue virions with deletions or defects in their genomes can be recovered from dengue patients. These defective viruses can only replicate with the assistance of fully functional viruses and they reduce the yield of the fully functional viruses. They are known as defective interfering (DI) particles. By administering additional, defined, DI particles to patients it may be possible to reduce the titre and duration of their viraemia. This, in turn may reduce the severity of the disease and the likelihood that dengue virus will be passed from the patient to a mosquito vector. This study estimates the number of DI particles that would need to be administered, and over what period, to have a significant effect on patient viraemia and subsequent dengue fever severity.

## Introduction

Dengue is caused by four serotypes (1-4) of a virus of the same name [1]. The viruses are transmitted between human hosts by Aedes mosquitoes, most commonly *Aedes aegypti*. Almost everyone living between the Tropics of Cancer and Capricorn are at risk of infection and an estimated 300 million infections occur each year [2, 3]. Disease symptoms range from a mild febrile illness to haemorrhagic fever and hypovolemic shock which, if untreated, is fatal in about 30% of cases [4]. Mosquito control programs have had little measurable effect on the number of reported cases of dengue [5], there is no vaccine and no disease specific therapy. Patients are treated by managing the symptoms with which they present.

Infection with one dengue virus (DENV) serotype probably results in life long immunity to re-infection with that DENV serotype but a second infection, with a different serotype, carries a significant risk of developing severe disease [6]. However, the onset of the severe symptoms in secondary infections usually occurs as the viraemia is waning and the secondary immune response is underway [7, 8]. There is a broad correlation between the magnitude of the viraemia in a dengue patient and the severity of the associated symptoms [9]. Any process that reduces the initial viraemia in dengue patients might reduce disease severity and also the risk that a mosquito feeding on the patient would become infected and pass the virus to a new host.

Populations of DENV include virions with genomes with defects ranging from single nucleotide changes [10] to deletion of more than 90 per cent of the genome [11]. Some of these are transmitted in nature for a year or more [10]. DENV virions containing genomes with extensive deletions interfere with the replication of wild type viruses. This phenomenon has been observed with a large number of viruses, mostly with RNA genomes [12, 13]. Furthermore, it has been possible to demonstrate that virions with defective genomes reduce the yield of virus from cells infected with wild type DENV and are known, therefore, as defective interfering (DI) particles [14–16].

There is an extensive literature on the activity of DI particles across a wide range of RNA viruses but interest waned in the 1990s [13, 17]. With the advent of tools to better define DI genomes and to produce artificial ones, there has been a renewed interest in their therapeutic potential and the possibility that they could be used to block transmission of agents such as DENV. However current mathematical models of dengue [18–20] cannot capture all the aspects of virus transmission and no model incorporates defective interfering (DI) particles. A few intracellular, intra-host and population models are available on different infectious diseases such as influenza, scabies, and optimal design for disease control [21–23]. This study uses data from 207 dengue patients in a real clinical setting [8] in order to estimate the therapeutic potential of DENV DI particles.

We propose a model inspired by the Clapham *et. al.* [19] and Frank [24] models. This model considers the antibody response in viral neutralization and the natural generation of DI particles. We build an ensemble, population, of models, in which each element in the population is a mathematical model with exactly the same framework, but where each model has a different set of parameter values for the same set of parameters. All of these values are calibrated in some appropriate way against multiple data [25]. In particular, we calibrate the data for plasma viral load and antibody response for 207 patients in our population of models (POMs). Most of the patients have high viraemia amplitudes during the illness. However, the antibody data has been collected on two random days within their febrile periods and that cannot explain the exact dynamics of the antibody, even asymptotically. With our POMs, we try to explore the range of variability in different cell-virus interactions and the immune responses. The POMs study is well-known in cardiac electrophysiology models [26, 27] but for infectious diseases, it is the first article to be reported for a large set of patient data. We develop a population of controls (POCs) to the population of symptomatic patients to attenuate the within-host viraemia level and reduce the days of febrile period. Specifically, bang-bang control is used to determine the minimum dose of DI particles which must be delivered to minimise the height and duration of the viraemia.

We propose that we can account for the inherent variability in the dengue infected patient data and find a modeling paradigm based on population of models and optimal control that allows us to quantify the effectiveness of DI particles in controlling the viraemia.

## Materials and methods

### Within-host viraemia dynamics

To explain the novelty of the present model, we must say that the competitive dynamics of the DI particles with virus is exhibited in the presence of the antibody response. While the model of Clapham et al. [19] included the role of antibody response in controlling the levels of viraemia, the model assumed that only standard virus is replicated within the host body. Defective interfering particles may also be responsible for the reduction in the production of standard virus [11, 14]. The dynamics of the present model is given by in the following set of ordinary differential equations

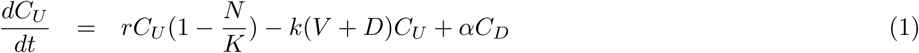

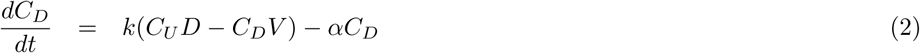

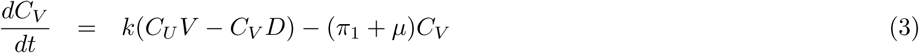

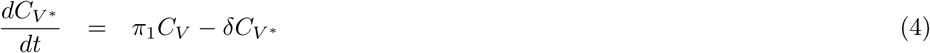

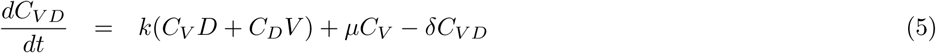

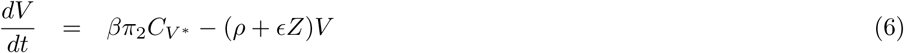

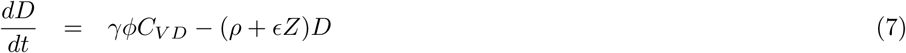

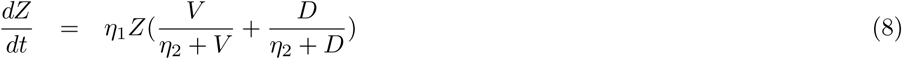

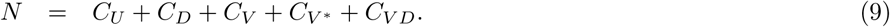

This new model describes the dynamics of standard virus (V) and DI particles (D) within the host. We consider the antibody response (Z) by the infected cells in virus neutralization. The present model is built with very specific aspects of dengue based on the models [17, 19, 24]. The uninfected target cells (*C_U_*) become infected and consequently produce four types of infected cells: infected by DI only (*C_D_*), virus only (*C_V_*), virus-infected and late enough for further infection (*C_V_ ∗*), and infected by both (*C_V D_*) (Fig 1). As the model of antibody dependent cell cytotoxicity (ADCC) is not as likely as virus opsonization, we do not consider ADCC in the present model. The assumptions that underpin our new model are described here. Bursting and cell lysis do not occur during the release of dengue virus particles. The infected cells are categorized in two classes according to their stages of infection: early and late. The early infected cells (*C_D_* and *C_V_*) are available for super-infection, but the late cells (*C_V ∗_* and *C_V D_*) are not. The immune response is strong in case of secondary infection leading to antibody-dependent enhancement (ADE) of the viraemia while it is very weak in case of primary infection. We consider the immune response in a simplified way such that the response is prominent only in the presence of significant antibody level, preferably in case of secondary infection. Both the defective and standard virus particles in this model are equally efficient in the competition of infection or replication.

**Fig 1.**
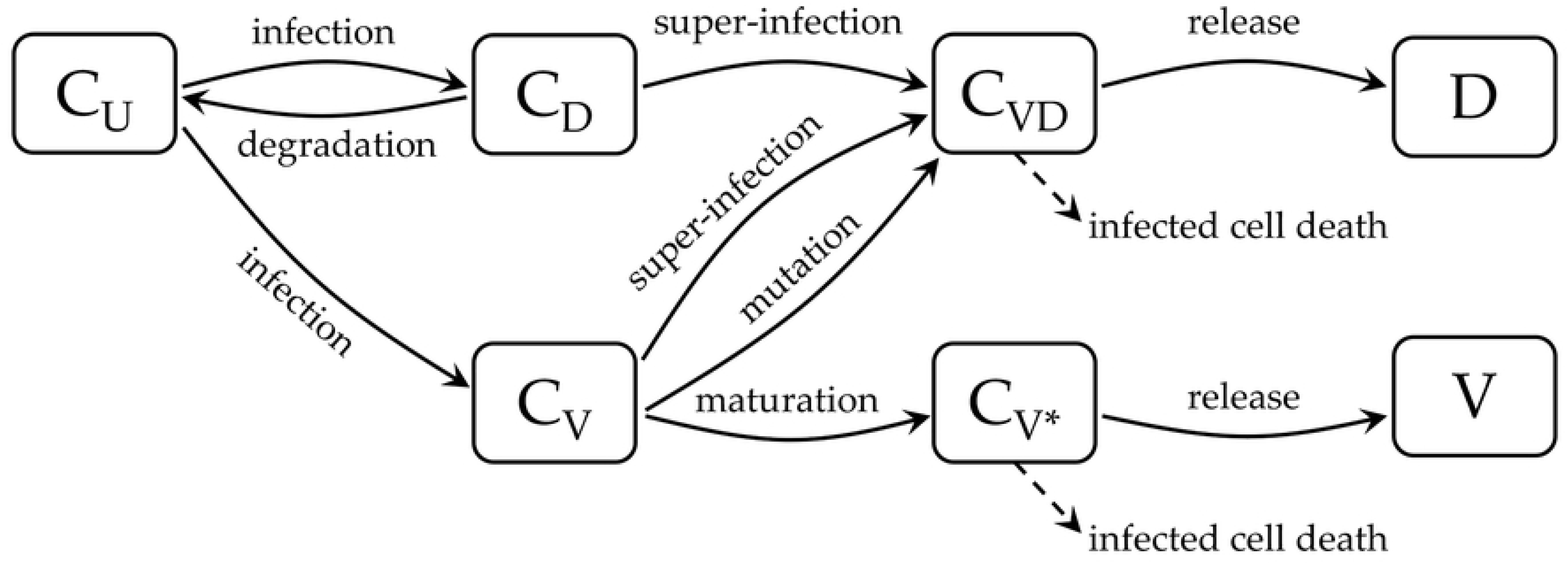
Schematic diagram of the within-host infection dynamics. *C_U_* denotes uninfected cells while *D* and *V* are the defective and infective virus particles, respectively. *C_D_*, *C_V_*, *C_V D_*, and *C_V ∗_* are the infected cells by *D*, *V*, both *D* and *V*, and infected by *V* only and matured, respectively.

Most of the model parameters must be estimated from the reported base values as the model is quite different from previous models, although the range of their values from the aforesaid papers [19, 24] are informative in creating the population of models. The initial conditions of *C_U_*, *C_D_*, *C_V_*, *C_V_ ∗*, *C_V D_* and *D* are considered constant at the start of infection. Only the initial viral load (*V*_0_) and antibody levels (*Z*_0_) for each patient have been sampled in the population of models. The patient specific parameters (*α, δ, η*_1_*, η*_2_*, π*_1_*, π*_2_*, φ*) are sampled using Latin Hypercube sampling (LHS) within the physiological range. LHS is a way of sampling high dimensional parameter spaces so that the number of samples does not scale with the dimension [28]. The way this is done is to discretise a *d* dimensional parameter space with some mesh and then place a cross in a box such that there is only ever one cross in each *d* − 1 dimensional subspace. A cross means that box is sampled at random for the *d* parameter values. The remaining parameters have been classified into two classes: natural human host parameters (*r* and *K*), which are constant in the complete POMs, and serotype-specific parameters (*β, ϵ, γ, k, µ, ρ*), which stay constant for a POMs of a particular serotype. We tabulate the description of the rate parameters in Table 1.

**Table 1.** Kinetic rate parameters used in the model.

| Parameters | Units | Descriptions | Source |
| --- | --- | --- | --- |
| Natural human host parameters |  |  |  |
| $K$ | cells per ml | Cellular carrying capacity of proliferation | [24] |
| $r$ | per day | Intrinsic rate of host cell proliferation | [24] |
| Serotype-specific parameters |  |  |  |
| $\beta$ | - | Number of $V$ released per $C_{V*}$ cells after packaging | [24] |
| $\epsilon$ | per day | Antibody mediated virus neutralization | [19] |
| $\gamma$ | - | Number of $D$ released per $C_{VD}$ cells after packaging | [24] |
| $k$ | per day | Rate of infection per virus | [19] |
| $\mu$ | per day | Mutation rate of $V$ to $D$ within host cells, turning $C_V$ cells into $C_{VD}$ cells | [24] |
| $\rho$ | per day | Natural clearance rate of $V$ and $D$ | [19] |
| Patient-specific parameters |  |  |  |
| $\alpha$ | per day | Rate of loss of DI particles within host cells, turning $C_D$ cells into $C_U$ cells | [24] |
| $\delta$ | per day | Death rate of infected cells | [19] |
| $\eta_1$ | per day | Proliferation rate of triggered immune response per infected cells by $V$ or $D$ | [19] |
| $\eta_2$ | - | Threshold parameter of the triggered immune cells proliferation | [19] |
| $\pi_1$ | per day | Rate of maturation of $C_V$ cells into $C_{V*}$ cells | [24] |
| $\pi_2$ | per day | Rate at which each $C_{V*}$ cells produces $V$ | [24] |
| $\phi$ | per day | Rate at which each $C_{VD}$ cells produces $D$ | [24] |
| $V_0$ | per ml | viraemia level on the day of infection | - |
| $Z_0$ | per ml | Level of immune response on the day of infection | - |

### Population of models

Variability inherently occurs in many biological and physiological measurements and we cannot avoid them. Every patient, for example, may have very different responses to an infection or a treatment and we need to account for this variability. Sometimes we aggregate the data and fit the model to the mean trajectory or choose a subset of the data as being representative or the hypothetically best sets of data and extrapolate those features to the large population. This can reduce the errors in measurement, but is unable to capture the intrinsic variability in the system. Hence, analysing models in a population from a set of measured data and exploring the hidden features intrinsic to the system is more effective for predicting physiological phenomena when there is inherent variability.

We use the Data2Dynamics package in Matlab for parameter estimation [29]. We generate multiple candidate models with parameters sampled by Latin Hypercube Sampling. We are at liberty to choose different criteria for our calibration. In original articles we calibrated to the range of the data [27], but this is somewhat crude. More recently, we proposed calibration based on matching the distributions in the data available [25]. This means that appropriate outputs from the POM matches the data in a distribution setting. In the present article we are following the earlier method as the amount of sampling is very large in the current system. In the first step of calibration, we try to calibrate to the median and other quartiles of the data but we can not fully capture the range of the data sets at the same time. For the whole data-set viraemia and antibody response, we estimate all the parameters and fixed the natural human host parameters (*r* and *K*) as constant and the remaining parameters are estimated again for the four different serotypes. The Latin Hypercube sampling is performed for each of the serotypes simultaneously with the serotype-specific parameters (*β, ϵ, γ, k, µ, ρ*) constant. From these population of models, we select only those models that cover the regions and range for all the biomarker results on each day of illness. We generate a very large initial POMs (20000) for each serotype and the calibrated POMs has been constructed by only those models that can capture the range.

### Optimal bang-bang control

There are two ways of implementing the control. One is continuous and differentiable. The other one is continuous but occurs as a step function and is known as bang-bang control, in which the control is either on or off. In practical settings bang-bang control is more appropriate for intervention and that is what we use here.

We follow the algorithmic steps for bang-bang control for a nonlinear system of ODEs as follows

1. Describe the system with the control variable as

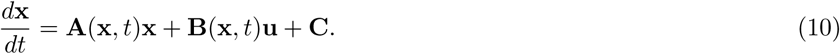
2. Construct the payoff functional in terms of running cost (*L*) and terminal cost (*φ*) functional as

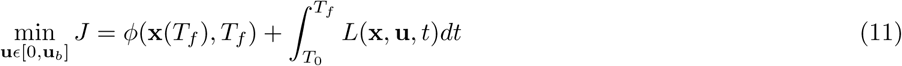

where **u** is the control variable, or vector of control variables, with bounds **0** ≤ **u** ≤ **u**_b_.
3. Construct the Hamiltonian

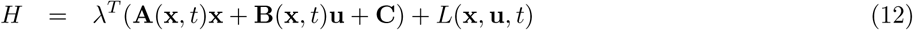

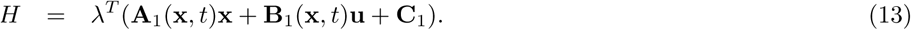

Here the lambda are the elements of the vector of Lagrange multipliers.
4. From the Pontryagin’s minimum principal [30] the switching function is

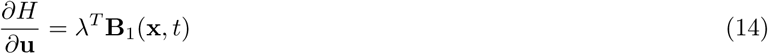

that determines the bang time (*τ*), when the control **u**(*t*) is on or off. The particular time points (*τ*’s) are known as the switching time points.
5. The optimal bang-bang control (**u**^*∗*^(*t*)) flips between the bounds, [0, **u**_*b*_] at the switching points as

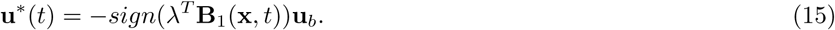

In the present study we use one control variable (*u*(*t*)), the administration of excess DI particles to the model to reduce the viral infection as well as quick clearance of the virus from the host. For the present POMs of four dengue serotypes, the range of the viraemia growth is large (approximately 10^3^ to 10^11^). For that reason it is difficult to decide on upper bounds of the control (*u_b_*) for these POMs. We determine the *u_b_* from the individual uncontrolled viraemia profile for each model considered to be controlled.

### Control strategy for dengue fever

As the objective to control dengue for the model within host, we construct an objective function in terms of the running cost functional only. The reason is that all the infection and virus naturally get cleared at the final time point and terminal cost is insignificant in such cases.

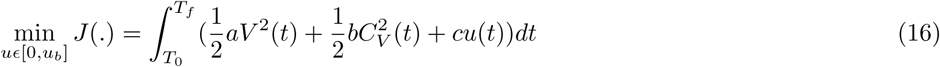

where *T*_0_ and *T_f_* are the initial and final time, and *a*, *b* and *c* are constants to be determined from the optimal control problem. In the course of control, we prefer to apply a bang-bang type control rather than a continuous control. Here, the administration dose rate (*u*(*t*)) of DI particles is the control variable. The medical nomenclature of the purified DI particles is therapeutic interfering particles or TIPs. In order to make the vaccination program cost-effective and reduce the time course of the vaccination process this information is included in the structure of the pay off function during the optimization. As the plasma viraemia (*V*) and the cellular infection of all kinds (*C_V_, C_V ∗_*) show a rapid growth in the first 2-4 days of the febrile period and are cleared within 10-12 days, we seek to minimize the peak of the viraemia (*V*) and virus infected cells (*C_V_*) that in consequence may help reduce all the infections. The DI particles within the host (*D*) compete with the virus for the uninfected cells (*C_U_*) and that is an advantage to introduce a large number of DI particles to inhibit the viral infection. The system of ODEs can be rewritten after introducing the control variable, *u*(*t*) as

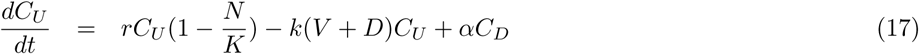

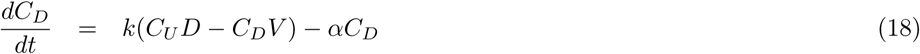

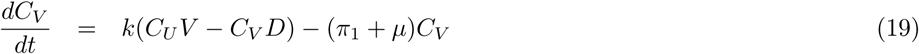

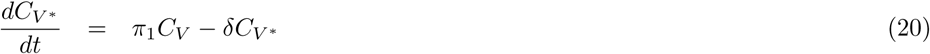

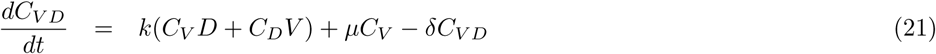

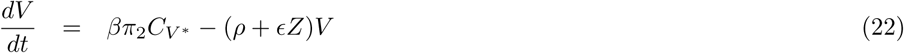

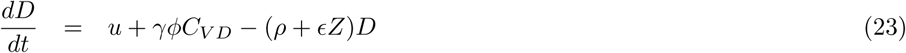

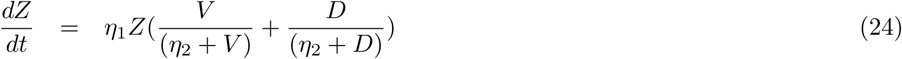

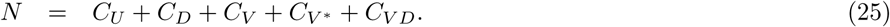

We assign bang-bang controls to the models from the POMs discussed above and obtain a population of controls (POCs) defined by the vectors of the amplitude of the bang of DI administration dose (*u*(*t*)) and on-off time duration (*τ*) of the bang-bang switches for the four serotypes.

## Results

### Population of models

From the experimental data, we have a set of 207 adult dengue patients with more than 3 days of fever [8]. Among them 38% and 40% of cases are DENV-1 and DENV-2 infections and a very low number of cases from DENV-3 (12%) and DENV-4 (11%). Most of the patients enrolled into hospital on days 2, 3 and 4 of their illness with high viraemia load in their blood samples. To build a model with an estimate of the day of infection using the day of illness is not appropriate. The days between the infection and start of illness are known as the incubation period for the plasma viraemia. For a large population of patients, it is difficult to frame the range of this time period in a dynamical model. To address this problem, we consider that the start of illness is a day in between the day of infection and maximum plasma viral load. The fever starts with a range of detectable viraemia load (*V*_0_) on the day the illness starts. Although DI particles are not observed directly in any prior study of blood viraemia trajectories they are known to occur naturally in viral infection systems. We may predict that from our POMs construction as they are generated naturally in viral infection systems. Fig 2 represents the calibrated POMs (black lines) with the reported plasma viraemia (red dots) for each of the four DENV serotypes for 10 days of their febrile periods. In the initial calibrated POMs, we found many viraemia models with large oscillations and abrupt growth in the antibody models. Although they satisfy the criteria to be included in the final POMs, they are omitted from the analysis as we cannot find any oscillatory behaviour in the reported viraemia data.

**Fig 2.**
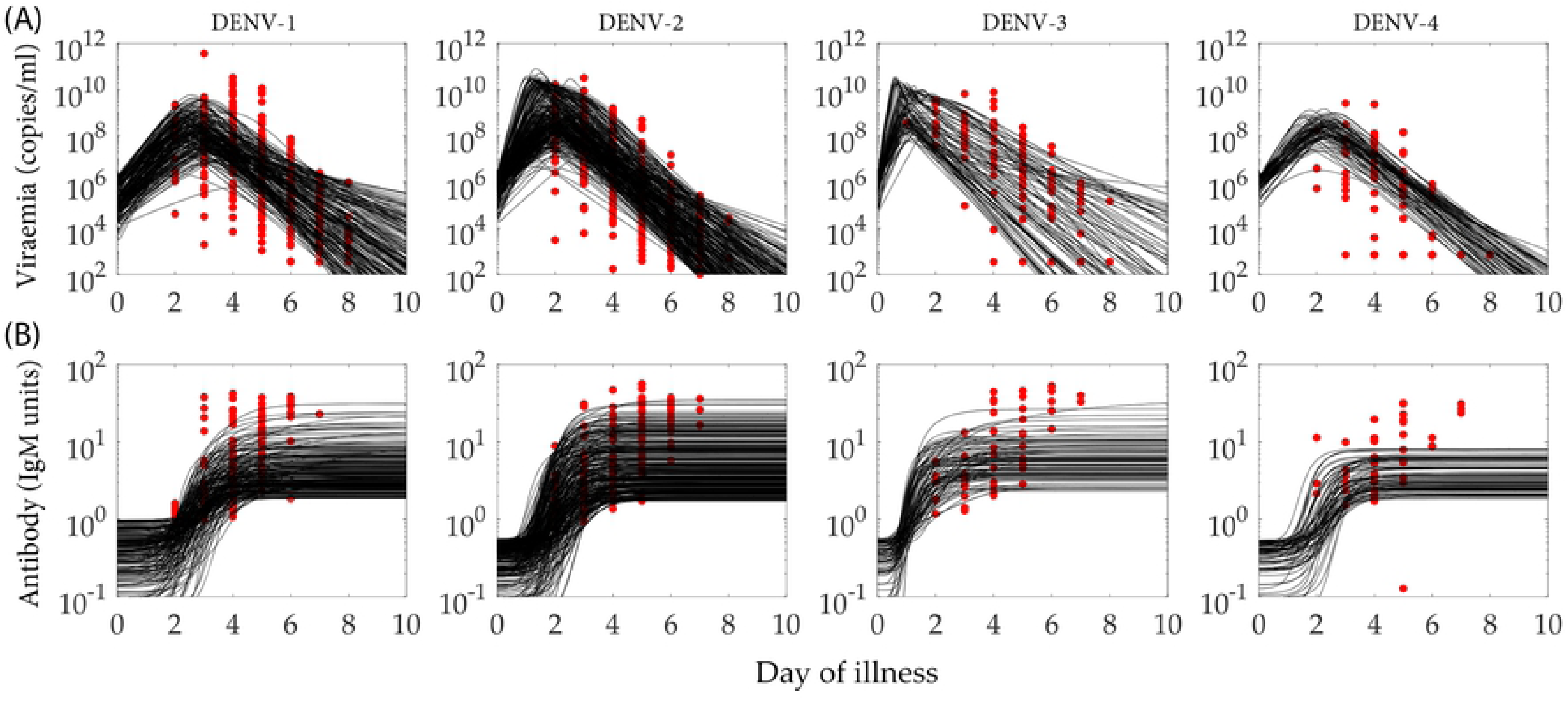
Population of models. (a) viraemia (V) and (b) antibody response (Z) obtained from clinical data (red) and calibrated models (black) included in the population for the four serotypes: DENV-1, DENV-2, DENV-3 and DENV-4. Both the biomarker levels are plotted with respect to the patient febrile time period.

In Fig 2, we present the POMs constructed (in black) based on the available biomarker data (in red). The data for the viraemia are regularly collected for every patients from day 2 to day 8 and that is reflected in the calibrated POMs nicely. But the available data for the antibody response is not that consistent as they appear randomly on any two of the days of illness. Calibration of the POMs for these data does not perform as effectively as for the viraemia population. To analyse the POMs for the four serotypes comparatively, we see that the POMs for DENV-2 is the most tightly calibrated with the biomarker data. The POMs for DENV-1 and DENV-4 are well calibrated in the dense region of the data and very few outlying data points cannot be captured in the POMs while DENV-3 POMs captures the spread of the data at every day of illness. In the case of DENV-2 and 3, the recurrence of tiny oscillations near the peaks of their rapid growths in the viraemia are more prominent than in DENV-1 and DENV-4 although that does not affect the antibody response. The antibody dynamics for the four serotypes are quite similar except in DENV-4. It is quite low in comparison to the other serotypes.

The spreads in different patient specific parameters for the four serotypes are shown in Fig 3. The rate of triggered immune response proliferation (*η*_1_) has notable differences in the case of DENV-4 from the other three serotypes. The effect of this narrow spread in *η*_1_ is reflected in the POMs for the DENV-4 antibody response in Fig 2B. The initial viraemia level (*V*_0_) spreads in a narrow domain for DENV-4 compared with the others and it makes the viraemia POMs in Fig 2A narrow. DENV-3, with its very narrow spread in *V*_0_, appears to be wide in the course of time. In all the cases, the low value of *η*_2_, the threshold of immune response proliferation, is inversely related to high level of *V*_0_.

**Fig 3.**
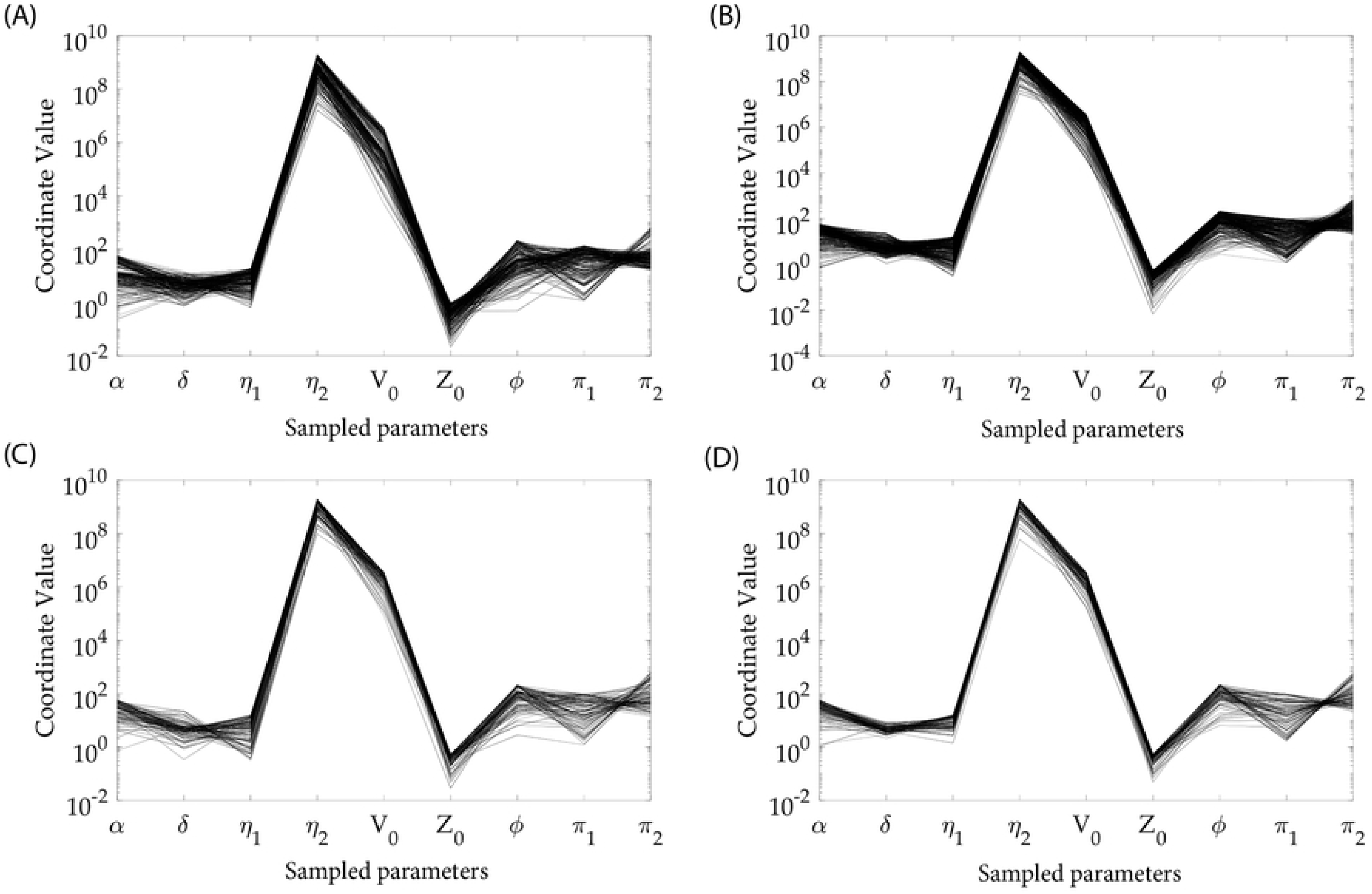
Variability in model parameters. Distribution of the patient specific model parameters plotted in parallel coordinates for (a) DENV-1, (b) DENV-2, (c) DENV-3, (d) DENV-4. The coordinate values (y-axes) are the values of the parameters while the parameters are noted in the x-axes.

In Fig 4, we depict the antibody response with respect to corresponding viraemia levels on every day of illness for further clarification of the calibration process. The black dots are the antibody-viraemia data points calculated from the accepted POMs on each day of illness. We show that most of the POMs results stay within the ranges of the biomarker data on day 3, 4, 5, 6 and 7 for all the four serotypes. On days 2 and 8, due to very low number of data-points, the range detection is not a reliable indicator of goodness of fit for the POMs..

**Fig 4.**
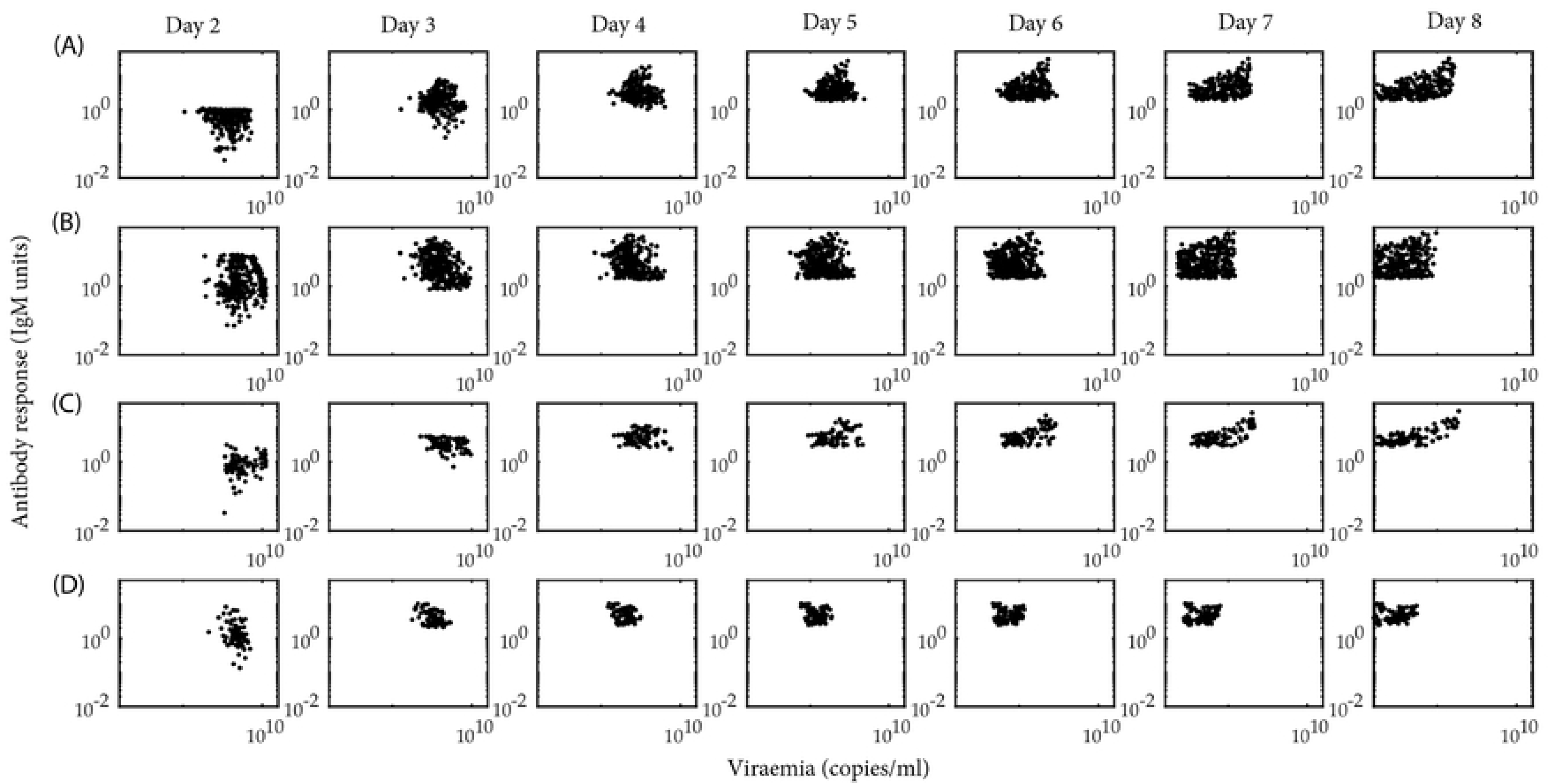
Relative variability in the biomarker data. Scattered phase plots of antibody response (Z) vs. the viraemia (V) on each day of illness for the dengue serotypes at each row: (a) DENV-1, (b) DENV-2, (c) DENV-3, (d) DENV-4.

For each of the patient-specific parameters, which have been allowed to vary in the population, the partial correlation coefficient (PCC) is calculated pairwise with the biomarkers calculated from the POMs. This correlation based approach can explore the sensitivity of the model parameters in association with the parameter variability. The PCC identifies one-to-one correlation between a particular parameter with the specific biomarker after removing the contributions of all the other variables. Thus it magnifies the one-to-one correlation between the parameter-biomarker pair. In Fig 5, we present three different heatmaps to quantitatively compare the PCC levels among the patient-specific parameters and the viraemia load, antibody response and accumulated DI particles levels across the four serotypes. Interestingly, although the POMs for viraemia load and antibody response show similar trends, the relation is not just straightforward if we look at the contributions of the model parameters through their PCC values.

**Fig 5.**
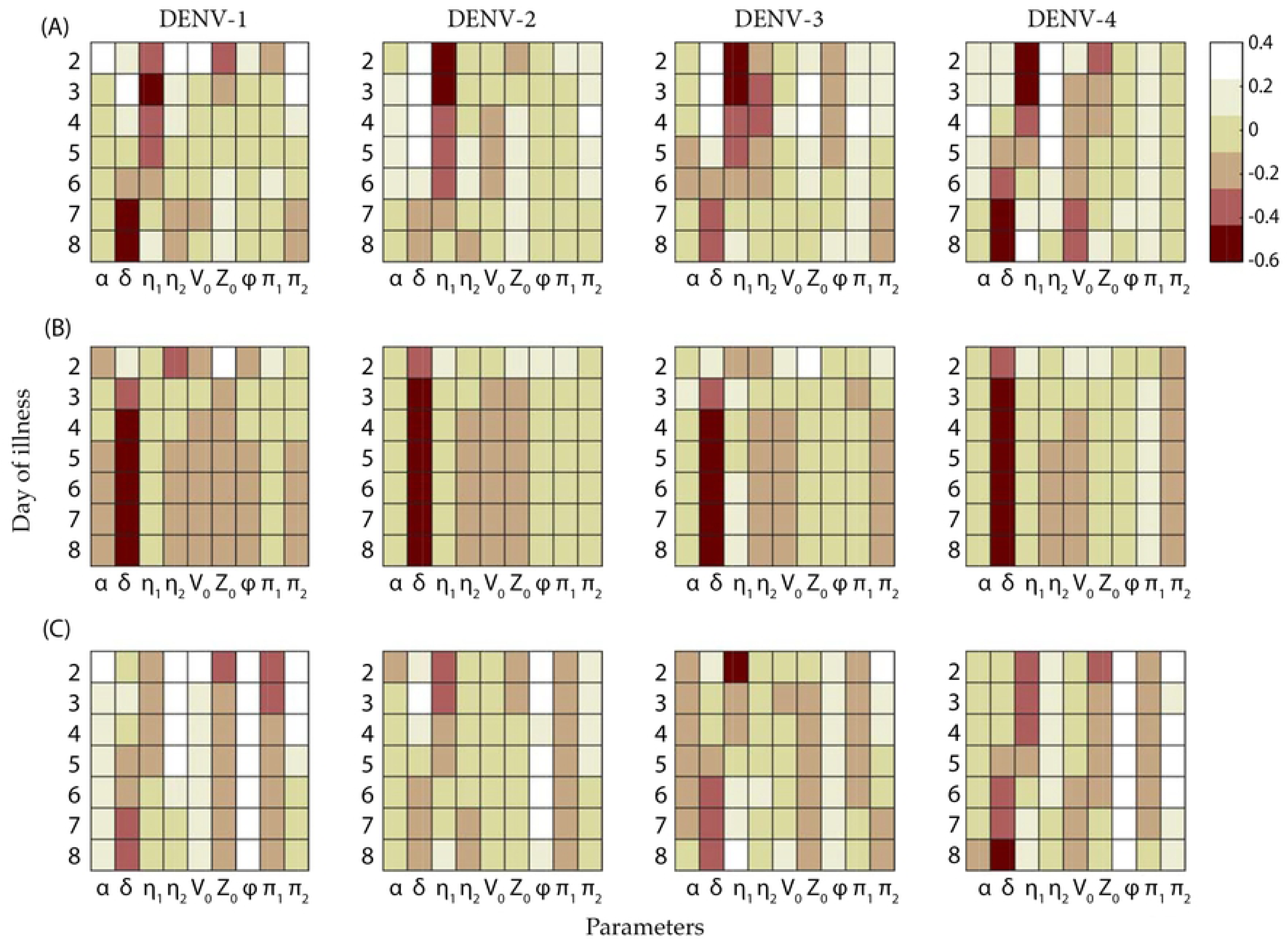
Partial correlation coefficient heatmaps. Partial correlation coefficient maps between the sampled model parameters and the model output of the biomarker levels: (a) plasma viraemia, (b) antibody level and (c) DI particles are calculated as a measure of sensitivity analysis for the four dengue serotypes.

In row Fig 5A, the PCCs of viraemia with different parameters are plotted. *δ* shows a transition from highly positive to highly negative correlation as long as the illness continues, while *η*_1_ goes in the opposite direction. However, *η*_2_ is not following a similar trend across the serotypes. To classify the PCCs for *η*_2_, DENV-1 and 3 are separable from the class of DENV-2 and 4. On the other hand, *α*, *V*_0_, *Z*_0_, *φ*, *π*_1_, *π*_2_ remain almost in the weak correlation regime with the viraemia for all the serotypes. In row Fig 5B, the PCCs of the antibody response with *δ* show high negative correlation while the rest of the parameter have no significant contributions. In the case of DI particles in row Fig 5C, all the parameters except *φ* and *π*_2_ appear with the same trend in Fig 5A, while *φ* and *π*_2_ show high positive correlation in all the serotypes on nearly every day of illness.

### Population of controls

Once the POMs have been constructed, we approach predicting the treatment for controlling the fever to the virtual population of dengue patient models. As the total number of qualified models in the POMs is large (221 for DENV-1, 306 for DENV-2, 93 for DENV-3, and 81 for DENV-4), we randomly choose 15% of the candidate models from the POMs of each serotype for the control experiment. During the random selection, we draw the models from the POMs with a uniform distribution and obtain 33,45, 13 and 12 models for DENV-1, DENV-2, DENV-3 and DENV-4, respectively. We could have chosen the best 15% of the best fitted models as the candidates for control experiment, but those do not appear in every domain of the POMs. In Fig 6, we present the viraemia, and DI particle levels before and after applying the control. For DENV-1, the viraemia lasts until day 10 keeping the control on for the whole period in most of the cases, while in case of the other serotypes the control shuts down approximately by day 8. The occurrence of the oscillatory peak in every few DENV-2 and DENV-3 models, pushes the control to higher dose although the viraemia cannot last beyond day 5.

**Fig 6.**
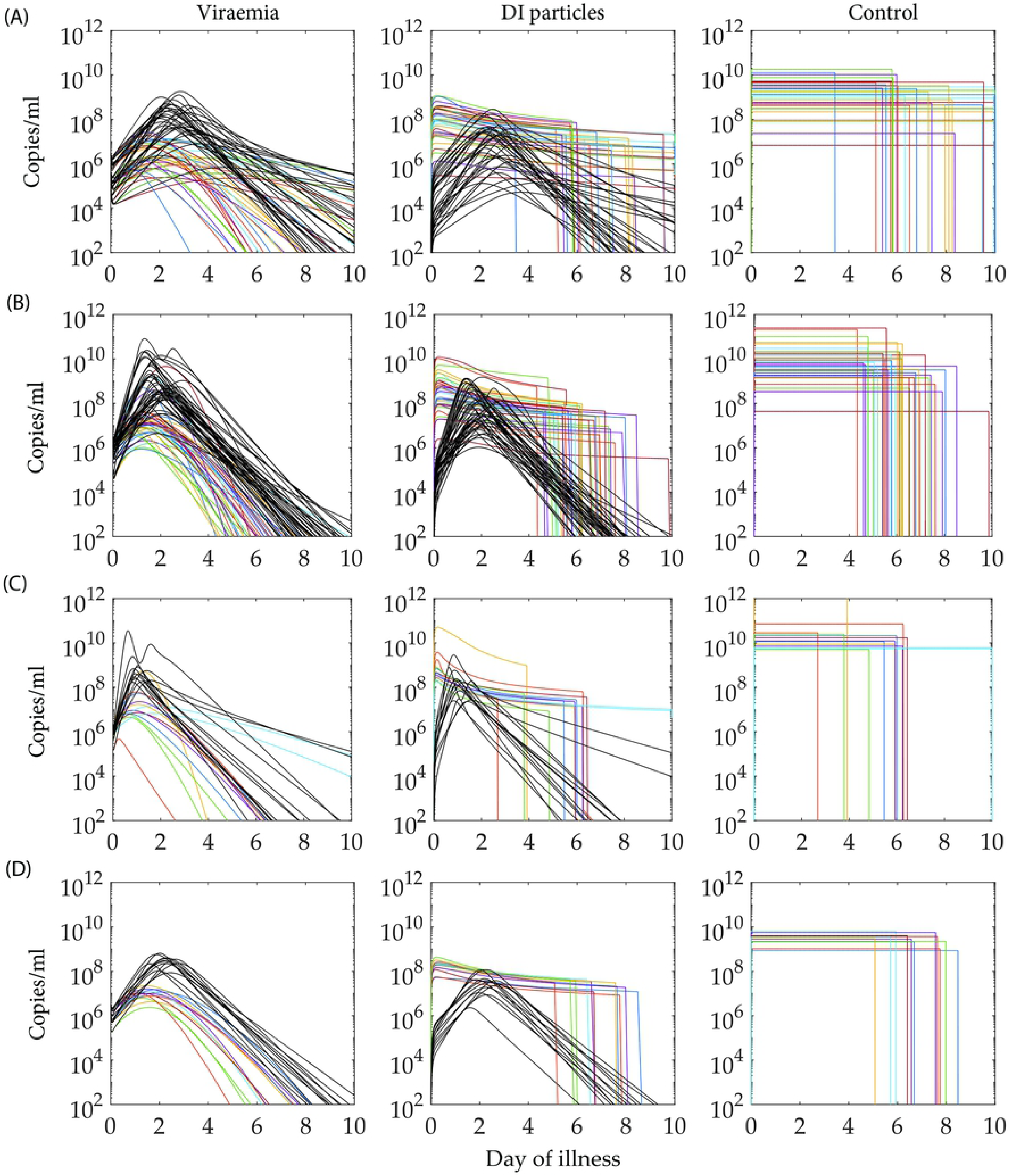
Effect of control in viraemia dynamics. The controlled viraemia, DI particles estimated from the optimal bang-bang control model and the control profiles for the dengue serotypes: (a) DENV-1, (b) DENV-2, (c) DENV-3, (d) DENV-4 are plotted in coloured lines while the black lines represent the corresponding profiles before applying the control.

The infected cellular dynamics also shows remarkable changes after the application of excess DI particles in the host system (Fig 7). The general trend before and after applying the control is observed in the *C_D_* cells, which is similar to that of the DI particles, as the DI particles are the major reason to generate the pool of *C_D_* cells. A similar relation is observed between the *C_V ∗_* cells with the viraemia profile as only *C_V ∗_* cells release potential virus into the body fluid. Interestingly, the application of the excess DI particles starts inhibiting both the virus and the *C_V ∗_* cells. The population of *C_D_* cells are produced from *C_U_* cells upon infected by *D* and *C_V D_* cells produce *D*. As a result, the pool of the DI particles drops sharply as soon as the control shuts down and the consequences are reflected in the *C_D_* and *C_V D_* cells.

**Fig 7.**
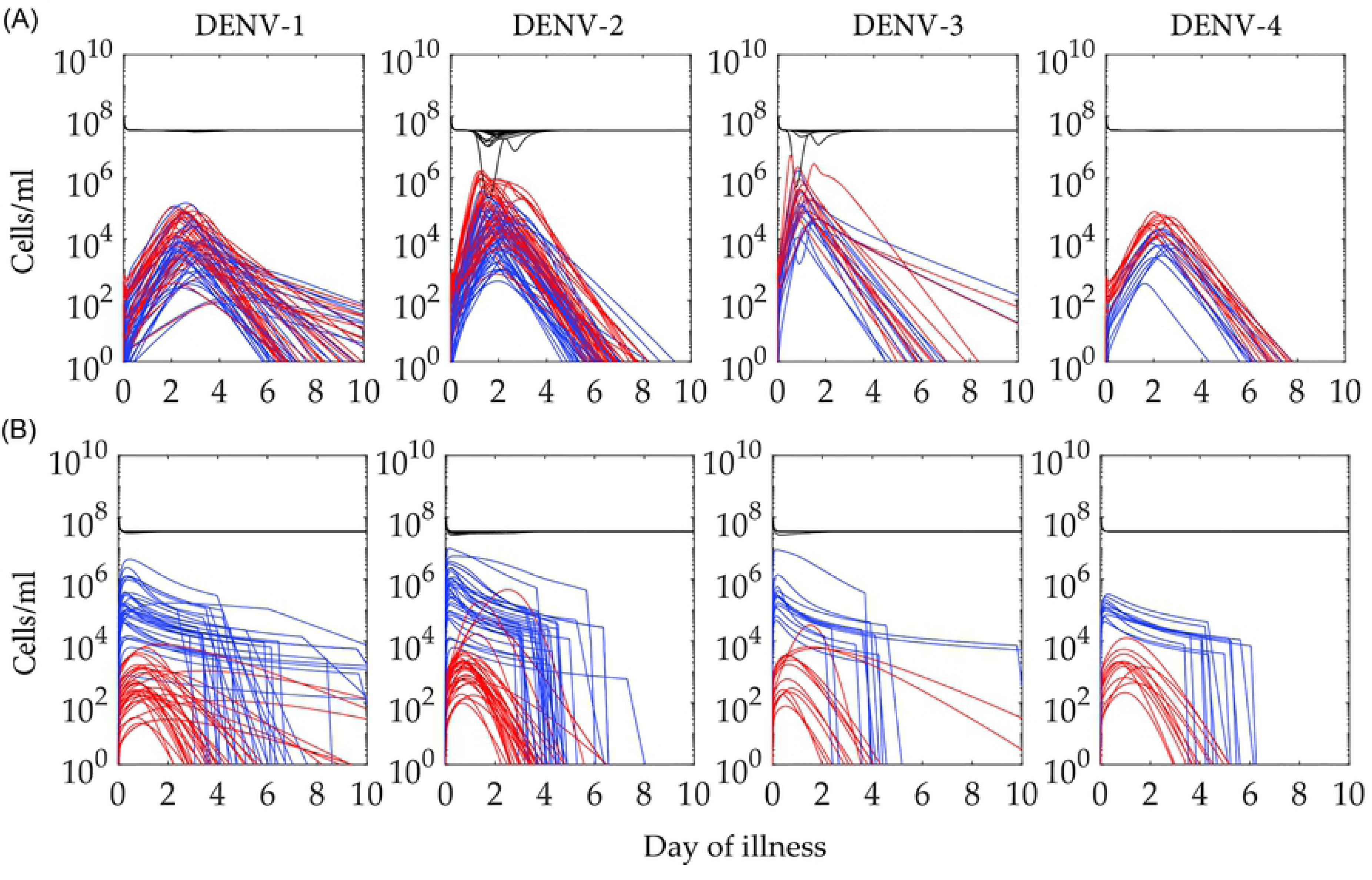
Effect of control in cellular dynamics. The dynamics of different cell types for the candidate models in the control experiment have been plotted during the febrile period. The uninfected cells (*C_U_*) (black), cells infected by DI particles only (*C_D_*) (blue) and cells infected by virus and are not available for superinfection (*C_V ∗_*) (red) are presented in black, blue and red lines, respectively in (a) before and (b) after applying the control, i.e., administration of excess DI particles.

If we consider the area under the control curve (*A*) as the cost of the vaccination, then an efficient control must be cost effective. To test the efficiency of the control, we estimate the area under the curve of the viraemia fold reduction (*R*) with respect to the area under the prescribed dose of control curve (*A*). Here the fold reduction (*R*) and control expense (*A*) are defined as

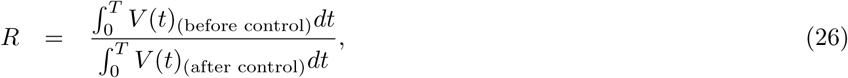

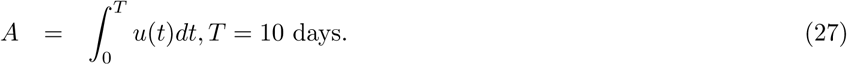

In Fig 8, we show the distribution of the viraemia fold reduction with respect to the control expense for all the four serotypes. Approximate monotonic increments are observed in *R*, with *S* for all the serotypes except DENV-2. For DENV-2, we find two separable clusters; one lies in the same cluster as the other serotypes and the other cluster appears with a completely opposite trend but at higher control expense.

**Fig 8.**
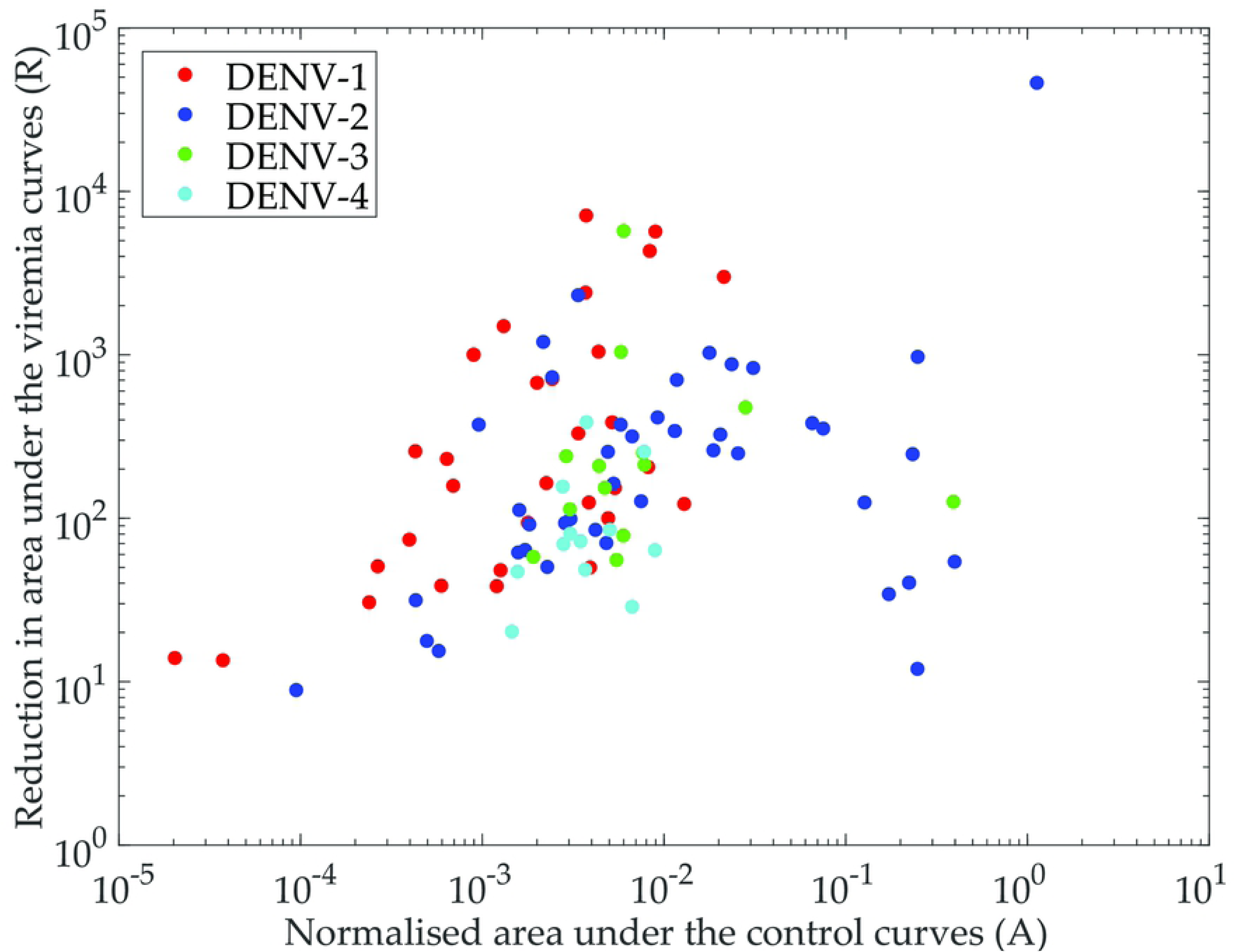
Control efficiency. The fold reduction in the viraemia is plotted with respect to the control, that is, the addition of excess DI particles for the four serotypes, DENV-1 (red), DENV-2 (blue), DENV-3 (green), DENV-4 (cyan).

## Discussion

The two prime interests of this paper are to capture the inherent variability in dengue infected patient data through a within-host model and predict efficient intervention to control dengue fever via administration of excess defective interfering particles (DIPs). We present the method of population of models (POMs) to execute the first goal and a population of bang-bang optimal control settings for the second aim. We show that the POMs not only capture the biomarker dataset but also provides the range of variability for each cell-virus interaction and its association with the biomarker kinetics in population and individual levels. A sub-population of the calibrated POMs are used with bang-bang control to reduce the viraemias in significant orders. In that case, the fever cannot reach severe dengue and the DI particles do not stop replicating. As per our findings, the antiviral property of the DI particles appears as a potential intervention strategy to attenuate the patient viraemia significantly.

We construct four serotype-specific populations of within-host models for dengue against the variability in the biomarker levels in blood samples of the admitted patients as reported [8]. The four POMs explore a range of patient-specific parameters, those in different combinations, produce four populations of feasible dengue models within the range of the experimental data. The calibration of the POMs helps us to discriminate and classify among the serotypes and inter-patient variability through the parameter variability and sensitivity. The aim of this methodology is not to look at the dynamics of isolated models in the population as any single model does not represent an individual. The aim is to incorporate variability in the same model and observe the whole population of patients with similar symptoms.

The variability appears in the population of the viraemia load and corresponding antibody response due to the differences in the patient-specific parameters. One of the crucial factors that drives this variability is the incubation period for an individual model. We want to mention that we trace the variability of incubation periods of an individual model in terms of the variability in viral load on day 0 of illness (*V*_0_) and that efficiently fits with the calibration process. The dynamics of the viraemia (*V*) is directly dependent on *δ*, *π*_1_, *π*_2_ for release after maturation of the infected *C_V_* to *C_V ∗_* and on the antibody response (*Z*) for clearance. Indirectly, the rate of infection (*k*) also drives the viraemia. Amongst these parameters, *δ* is in strong positive correlation with *V*, *Z* and *D* and that gradually leads to a flip as the viraemia dies with the days of illness, but *π*_1_ is weakly correlated all the time. The variability of highly correlated parameters stay within a narrow range and calibrates tightly with the biomarker data, but weakly correlated parameters spread over wide ranges to generate models with similar behavior (Fig 3 and 5).

In the Ben-Shachar *et. al.* [31] statistical model, the populations of infected patients have been classified according to the disease severity across the serotypes and the variability in their immune responses. Although this study is more concerned with the immune response, they predicted the relation among virus replication rates with the timing of the viraemia peaks over the days of illness. Our POMs results show consistency with their observations when we demonstrate the variability for different parameters. DENV-1 and DENV-4 reach the viraemia peaks after the symptom onset, while the peaks appear before the onset of the symptoms in case of DENV-2 and DENV-3 and it depends on the degree of infection (Fig 2). Although, the few relatively high peak heights in viraemia data for DENV-1 cannot be captured in our model.

Among the reported infections of the hospitalized patients in our model, most of the DENV-1 infected patients have primary infection while the majority of the patients with the other serotypes are reported as secondary infection. A careful observation of the POMs of the viraemia profiles enables us to find the growth rate of the viraemia for most of the models in, with the DENV-2 and DENV-3 POMs growing faster than the others. We explain this rapid growth in terms of the antibody dependent enhancement (ADE) that only occurs in secondary infection [6]. In case of primary infection, the immune response is triggered very slowly and the viraemia is almost cleared when the response level is significant. On the other hand, the same response for the secondary infection is very rapid and prominent.

In the articles of Clapham *et. al.*, two different within-host models for dengue infection have been presented for DENV-1 and DENV-2. They found variability in the rate of infection (*k*) only and that was used to discriminate between the ranges of viraemia loads [20]. Later they have fitted another model with a direct and indirect effect of the antibody response through free virus neutralization and infected cell death [19]. In this present article, we keep *k* and *ϵ* constant for each serotype and included only the direct antibody response for virus particles (standard or defective) neutralization and the antibody response is triggered by both of the free virus and free defective particles. The variability in the antibody response is captured by *η*_1_ and *η*_2_ and their contributions are reflected in the POMs. The greater the proliferation (*η*_1_) rate varies, the more the antibody plateau widens (Fig 2 and 3). Notably, in the case of DENV-4, the spread for both of *η*_1_ and *η*_2_ are narrow. Again, the strong negative correlation of *η*_1_ with the viraemia does not appear to be significant in comparison with the case of the DI particles and *Z*. This may explain the intensity of the triggered antibody response being more effective on *V* than *D*.

Another significant outcome of such a population level modeling approach is in the quantitative prediction of vaccination or any kind of intervention strategy. We use an optimal bang-bang control approach to add excess DI particles in to the system to reduce the viraemia. Previously, Rodrigues *et. al.* showed optimal control for dengue using vaccination compartment inside an epidemic viewpoint [18]. But intervening individual human host models within a population has not been observed yet. Furthermore, the naturally occurring defective interfering particles have not been utilized in dengue control before.

Bang-bang control is a prominent optimization tool in dynamical programming for linear systems and can be solved easily using boundary value problem (BVP) solvers [32]. But a nonlinear two-point boundary value problem (TPBVP) such as our present model cannot be solved directly with traditional solvers. We use the forward backward sweep method, where ODE solvers are used twice: forward for the state equations and backward for the costate equations [33]. Then we update the switching function(*∂H/∂u*) and control (*u*(*t*)) [34, 35]. In most of the cases, for such nonlinear models, nonlinear programming is mostly used for calculating discontinuous controls. We use the same Pontryagin’s minimum principle and solve the discontinuous right hand side of the state and co-state equations. We note that this method needs many more iterations than continuous control methods to converge. However, for models with strong non-linearity such as stiff and oscillatory control problems, this method is reasonably efficient.

We perform the control experiment on a randomly chosen 15 per cent of models from the calibrated population of models for each serotype. In Fig 6, the population of controls (POCs) profiles for the four serotypes are quite self-explanatory. As the replication of the DI particles depend on the replicative machinery of the standard virus, the excess DI particles are rapidly cleared out of the host system as soon as the control shuts down and viraemia is cleared. We ensure the amplitude of the control, i.e., addition of excess DI particles, to be equivalent to the level of viraemia peak during computing the controls, otherwise the amount of the DI particles are not sufficient to reduce the viraemia peak. Our aim is to keep the viral load approximately below 10^8^ but for DENV-2 and DENV-3 it is difficult to achieve that even after applying 10^11^ of DI particles. The reason behind this is the higher rates of virus replication (*β* and *π*_2_) in DENV-2 and DENV-3 as mentioned before. In the cases of DENV-1 and DENV-4, as soon as the DI particles start boosting, the viral load drops quickly, as DI particles interfere in the virus replication. Very tiny persistent oscillations in the case of DENV-2 and DENV-3 in all the cell types and viraemia also validates the same conclusions.

To examine the efficiency of the control experiment, we refer to the scatter plot in Fig 8 for the measured control expense (A) and the corresponding reduction in viraemia (*R*). For DENV-1, DENV-3 and DENV-4, most of the models are in the left half of the figure (i.e., *A ≤* 10^3^) while DENV-2 has many more models in the high *A* domain (i.e., *A ≥* 10^3^). In most of the cases for DENV-1, the reduction (*R*) is higher than the other serotypes at low expense on control (*A*) and that makes the control for DENV-1 as the most efficient. The present model predicts that large numbers of DI particles would be administered to DENV patients to have any effect on viraemia as patients only become symptomatic and seek medical assistance at the time of peak viraemia or soon after. The model also assumes that DI particles and wild type viruses are of equal fitness when competing for replicative machinery within host cell. If, however, DI particles are interfering with replication of wild type viruses by enhancing production of interferon or some other mediator, then a single DI particle/genome may elicit a response in the host cell that interferes with the replication of large number of wild type viruses. In addition, there exists no specific metric that may provide room to define the efficiency of the DI particles. A distribution of DI particle with variability in their competitions with the virus particles for the replication and packaging can be modeled to predict the efficiency of the DI particles through successive passages. Existing models and experiments with DI particles assume the efficiency of the DI particles inversely proportional to their nucleotide lengths though the nucleotide lengths cannot decide on DIP efficiency. A single cell stochastic model with distribution of DIPs and their evolutionary aspects may open a new avenue to explore the DIP efficiency.

Despite the availability of real clinical data for the admitted patients and experimental success, the intra-host dengue virus dynamics is not explored well. As a consequence, the virus transmission dynamics to mosquitoes is not clear. This paper explores the variability regime of the intra-host DENV dynamics across a population of patients for the four DENV serotypes. These POMs are able to predict the effective roles of the virus replication and subsequent immune response to determine the within-host viraemia characteristics. For the same patients population, a human to mosquito transmission model is underway. Those results may explore the quantitative analysis of infected patients turned into infectious and their infectiousness in terms of the transmission. Addition of minimal amount of defective particles leads to significant reduction in the viraemia characteristics reflecting the potential anti-viral property to be manifested in dengue control.

## Acknowledgments

We thank Nguyen *et. al.* for providing the data collected at the Hospital for Tropical Diseases, HCMC, Vietnam. This work is supported by a DARPA INTERCEPT Program (HR0011-17-2-0036). We thank all those engaged in the DARPA INTERCEPT Dengue TIPs for suggestions and discussions. We thank the High Performance Computing (HPC) and Research Support of QUT for the computational facility.

## Author Contributions

**Conceptualization:** TM, SC, JA, KB.

**Data curation:** TM, SC.

**Funding:** JA, KB.

**Investigation** TM, JA, KB.

**Methodology** TM, KB.

**Writing - original draft:** TM, KB.

**Writing - review and editing:** TM, SC, JA, KB.

